# Viral protein engagement of GBF1 induces host cell vulnerability through synthetic lethality

**DOI:** 10.1101/2020.10.12.336487

**Authors:** Arti T Navare, Fred D Mast, Jean Paul Olivier, Thierry Bertomeu, Maxwell Neal, Lindsay N Carpp, Alexis Kaushansky, Jasmin Coulombe-Huntington, Mike Tyers, John D Aitchison

## Abstract

Viruses co-opt host proteins to carry out their lifecycle. Repurposed host proteins may thus become functionally compromised; a situation analogous to a loss-of-function mutation. We term such host proteins viral-induced hypomorphs. Cells bearing cancer driver loss-of-function mutations have successfully been targeted with drugs perturbing proteins encoded by the synthetic lethal partners of cancer-specific mutations. Synthetic lethal interactions of viral-induced hypomorphs have the potential to be similarly targeted for the development of host-based antiviral therapeutics. Here, we use GBF1, which supports the infection of many RNA viruses, as a proof-of-concept. GBF1 becomes a hypomorph upon interaction with the poliovirus protein 3A. Screening for synthetic lethal partners of GBF1 revealed ARF1 as the top hit, disruption of which, selectively killed cells that synthesize poliovirus 3A. Thus, viral protein interactions can induce hypomorphs that render host cells vulnerable to perturbations that leave uninfected cells intact. Exploiting viral-induced vulnerabilities could lead to broad-spectrum antivirals for many viruses, including SARS-CoV-2.

**Summary:** Using a viral-induced hypomorph of GBF1, Navare et al., demonstrate that the principle of synthetic lethality is a mechanism to selectively kill virus-infected cells.

## Introduction

RNA viruses are prevalent and pervasive pathogens responsible for many global health crises, including the COVID-19 pandemic (Carrasco-Hernandez et al., 2017; Enard and Petrov, 2020; Rosenberg, 2015; Woolhouse and Gaunt, 2007). RNA viruses typically have high mutation rates enabling them to rapidly adapt to new cell types, infect new host species, evade host immune responses, and quickly develop antiviral drug resistance (Sanjuán et al., 2010). Currently, the repertoire of U.S. Food and Drug Administration approved antivirals is limited, targeting only eight out of the known 214 human-infecting RNA viruses (Heaton, 2019). Almost exclusively, these U.S. Food and Drug Administration approved drugs are all designed to target viral proteins (Heaton, 2019; Woolhouse and Brierley, 2018). The dearth of antivirals is likely impacted by the many challenges faced in antiviral drug development, including the small list of target viral proteins due to their compact genomes, the quick emergence of escape mutants due to the high mutation rates prevalent in many RNA viruses, and the limited therapeutic range of antivirals due to the diversity of RNA virus serotypes which, akin to antimicrobial resistance, has led to limited strain-specific use, or even discontinuation of use, for many antivirals (Heaton, 2019; Irwin et al., 2016; Pennings, 2013; van der Vries et al., 2013). Yet, the same compact-sized genome that gives these viruses an edge over antivirals also makes them obligatory pathogens that rely on host proteins for survival. Thus, although viral genomes drift, they often maintain reliance on the same subset of host factors (Gordon et al., 2020a; Heaton, 2019). The idea of exploiting this over-reliance on their host to develop host-directed antivirals to interfere with host cell factors that are required by the virus, or to more broadly influence immune responses is gaining traction (Gordon et al., 2020b; Kaufmann et al., 2018; Mast et al., 2020; Prussia et al., 2011).

Host-based therapies expand opportunities for treating viral infections (Brass et al., 2008; Krishnan and Garcia-Blanco, 2014; Zhou et al., 2008). To proliferate, viruses must co-opt common host pathways and cellular machineries by forming protein-protein interactions with host proteins (Basler et al., 2019; Carpp et al., 2014; Gordon et al., 2020b; Lum and Cristea, 2016; Saeed et al., 2020; Stukalov et al., 2020). Due to the essential requirement of these interactions, they are likely to be conserved within viral lineages and can be targeted for developing broad-spectrum therapeutics (de Chassey et al., 2014; Meyniel-Schicklin et al., 2012; Pfefferle et al., 2011). Recently, an analysis of published virus-host interaction datasets revealed RNA viruses frequently engage so-called “multifunctional host proteins”, a set of 282 highly connected proteins as assessed by their protein-protein interaction networks (Heaton, 2019; Navratil et al., 2009). Targeting these multifunctional host proteins with drugs, many approved by the U.S. Food and Drug Administration could disrupt several crucial steps of viral replication. Inhibiting a virus by targeting the host reduces the possibility of drug resistance and is potentially broad-spectrum if more than one virus relies on the same host protein. However, inhibiting these multifunctional proteins by drugs may also elicit adverse effects on the host because often these proteins tend to be essential and serve as hubs of complex protein interaction networks (Heaton, 2019; Zotenko et al., 2008). Thus, host-based therapeutic targets should be chosen carefully to avoid potential serious adverse side effects and, ideally, strategies that selectively affect only infected cells should be sought.

The principle of synthetic lethality offers an opportunity for selectively targeting virus infected cells by drugging synthetic lethal (SL) interactors of virus-targeted multifunctional protein hubs (Mast et al., 2020). Synthetic lethality occurs between two genes when a loss-of-function mutation in either gene has little impact on cell viability, but becomes detrimental when paired together resulting in cell death (Dobzhansky, 1946; Hartwell et al., 1997) (Fig. 1A). Such lethal genetic combinations, known as “synthetic lethal pairs” (Nijman, 2011a), are one of many forms of genetic interactions that can occur within cells (Boone et al., 2007; Dixon et al., 2009; Drees et al., 2005; Horlbeck et al., 2018). The existence of synthetic lethality reveals important aspects of the genetic architecture of cells, demonstrating the presence of genetic buffering in organisms due to functional redundancy (Horlbeck et al., 2018; McManus et al., 2009). This SL concept has been successfully applied to cancer therapy and host-targeted drug development (Farmer et al., 2005; Kaelin, 2005; Mendes-Pereira et al., 2009; Turner et al., 2008; Wiltshire et al., 2010) (Fig. 1B). For example, loss-of-function mutations in the DNA repair genes encoded by breast cancer type 1 and 2, *BRCA1* and *BRCA2*, cause breast and ovarian cancer but exhibit enhanced sensitivity to inhibitors of poly ADP-ribose polymerase (*PARP*), another DNA repair enzyme (Farmer et al., 2005). PARP inhibitors selectively killed cancerous cells carrying the loss-of-function *BRCA* mutation while sparing noncancerous cells (Bryant et al., 2005) and in a clinical trial, PARP anticancer drugs showed a significantly longer progression-free period in patients with breast cancer (Litton et al., 2018). Synthetic lethality-inspired anticancer therapy provides avenues for improved drug specificity and efficacy at lower doses, thereby limiting side effects (Beijersbergen et al., 2017). Here, we extend the application of this synthetic lethality principle to host-derived antiviral targets.

**Figure 1.**
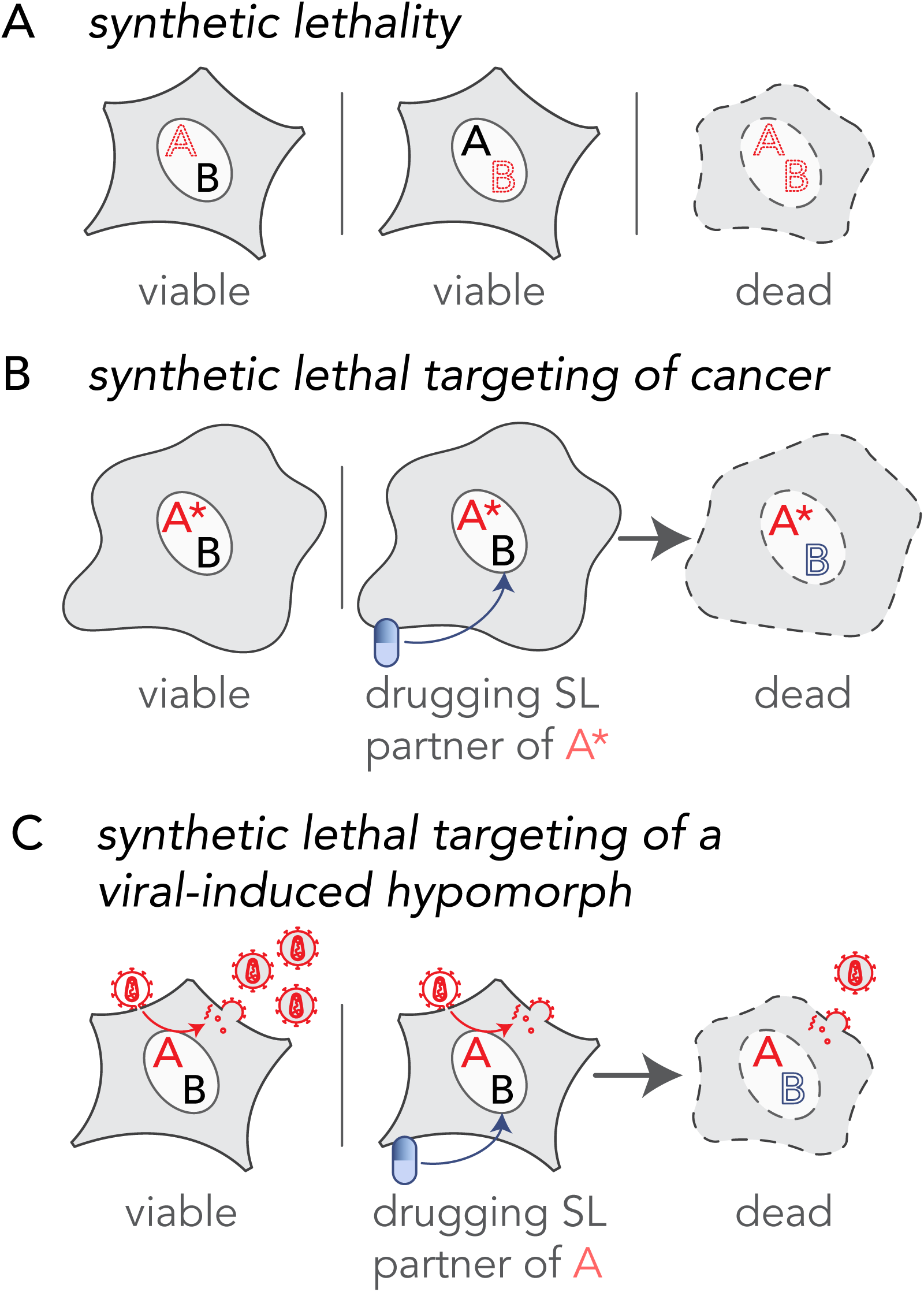
Extending the principle of synthetic lethal interactions to a virus-induced hypomorph. (**A**) Synthetic lethality is an extreme negative genetic interaction occurring between two genes. Here, genes ‘A’ and ‘B’ are not essential, and the cell remains viable upon the loss of either gene, depicted by red dotted outline of ‘A’ or ‘B’, individually. However, when these deletions are combined in a single cell, as visualized in the third panel, this double loss of function critically impairs the cell, resulting in its death. Such gene-gene combinations are termed synthetic lethal (SL) partners. (**B**) The principle of synthetic lethality has been successfully exploited in the development of certain cancer therapies by targeting the synthetic lethal partner of the cancer-causing oncogene, depicted by a red ‘A’. In the cancerous cell, gene ‘A’ has been mutated, depicted as ‘A*’, leading to an enhanced dependency by the cancer cell for its synthetic lethal partner ‘B’. Drugs that target the otherwise nonessential gene B induce cell death when combined with its SL partner, A*. Therefore, inhibiting the function of B can selectively kill cancerous cells while sparing noncancerous bystander cells. (**C**) Like the example in cancer (B), a viral infection provides opportunities for specifically targeting infected cells by synthetic lethality. When a cell is infected, host factors, depicted as the red letter ‘A’, are recruited by viral proteins to support viral reproduction. The normal function of the host factor is thus attenuated by the presence of the virus, inducing a hypomorph, red letter ‘A’, which sensitizes the infected cell to inhibition of its synthetic lethal partner by an inhibitory drug.

Virus infection is a perturbation of the host protein-protein interaction (PPI) network. Interactions between virus and host proteins usurp normal protein functions and rewire host PPI networks. Host proteins are considered proviral if loss of function renders the host cell resistant to infection, and antiviral if loss of function improves cell permissibility to infection. Infected cells exhibit altered metabolic requirements (Thaker et al., 2019), signaling pathways (Gaur et al., 2011), intracellular transport pathways (Belov et al., 2007) and other morphological and molecular characteristics relative to the noninfected cells. In such situations, infected cells may depend on a different complement of proteins than their uninfected counterparts (Mast et al., 2020). This state-specific vulnerability may be a target for host-based therapeutics based on the well-established principle of synthetic lethality. For example, if two host cell proteins have a SL relationship and the function of one protein is hijacked by a viral protein, then cells may become dependent on the function of the second protein. In contrast, cells that are not altered by the virus, i.e., those that are uninfected, will be unimpacted by blocking the second protein, since the elimination of a single half of the SL pair does not result in a phenotype. Rational targeting of SL protein pairs in which the function of one partner is reduced specifically in the infected cell; a situation equivalent to the mutant gene in cancer, is a novel framework for taking advantage of the intrinsic differences of infected cells to achieve selective targeting (Fig. 1C). We hypothesize that viral-host PPIs generate protein-based, viral-induced (vi)-hypomorphs of host factors in infected cells, thereby specifically sensitizing infected cells to targeting SL/synthetic sick partners of these vi-hypomorphs. To test this hypothesis, we selected the Golgi-specific brefeldin A-resistance guanine nucleotide exchange factor (GBF1) (Claude et al., 1999) because it is a critical proviral host factor for the replication of several families of RNA viruses, including *Picornaviridae, Coronaviridae, Flaviviridae, Herpesviridae, Filoviridae*, and *Rioviridae* (Belov et al., 2008b; Carpp et al., 2014; Farhat et al., 2018; Goueslain et al., 2010; Lanke et al., 2009; Martínez et al., 2019; Verheije et al., 2008; Yamayoshi et al., 2010). Considering so many viruses rely on GBF1, it seems unlikely that these viruses would readily overcome GBF1 dependence. GBF1 mediates recruitment of coat proteins (Manolea et al., 2008), lipid modifying enzymes (Ellong et al., 2011), and protein tethers (García-Mata and Sztul, 2003) to Golgi membranes. It thus plays a central role in vesicular transport through the Golgi, the structural integrity of Golgi membranes, and the maintenance of lipid homeostasis (Beller et al., 2008; Donaldson and Jackson, 2011; Guo et al., 2008; Sáenz et al., 2009; Soni et al., 2009). With its dynamic membrane-modulating functions GBF1 has also been implicated in coatomer-dependent protein delivery to lipid droplets (Kaczmarek et al., 2017; Soni et al., 2009).

Many RNA viruses encode proteins that bind GBF1 directly, including the nonstructural proteins 3A of poliovirus (Belov et al., 2008a; Teterina et al., 2011) and coxsackievirus (Wessels et al., 2006a; c), and nonstructural protein 5 of dengue (Carpp et al., 2014). Recently, two SARS-CoV-2 proteins, membrane (M) and orf6, were identified to be directly binding or in close proximity of GBF1, respectively (Laurent et al., 2020; Stukalov et al., 2020). In the case of poliovirus infection, 3A redistributes GBF1 to viral replication complexes during early stages of replication and subverts its guanine nucleotide exchange factor (GEF) function in the infected cells (Carpp et al., 2014; Richards et al., 2014; Wessels et al., 2007, 2006b; c), suggesting that poliovirus protein 3A may attenuate GBF1’s normal function creating a hypomorph, rendering cells susceptible to disruption of proteins synthetically lethal with *GBF1*. Here, we provide proof-of-concept that SL partners of vi-hypomorphs can be targeted to selectively eliminate infected cells while leaving uninfected cells intact. We do this by performing a genome-wide chemogenomic CRISPR screen to identify SL partners of *GBF1*, validating the top candidates, and demonstrating that shRNA-mediated silencing of the *GBF1* SL interacting partner, *ARF1*, selectively kills cells expressing poliovirus protein 3A.

## Results and Discussion

In order to identify putative synthetic lethal partners of *GBF1*, we screened a high-complexity extended-knockout CRISPR library of 278K single guide RNAs (sgRNAs) that target 19,084 RefSeq genes, 20,852 alternatively-spliced genes, and 3,872 predicted genes, among additional controls, in NALM-6 human B cell precursor leukemia cells (Bertomeu et al., 2018) (Fig. 2A). These cells harbor a genomic doxycycline-inducible Cas9 that enables regulatable, uniform, and robust gene silencing across the pooled library (Wang et al., 2014). Relative changes in sgRNA frequencies were obtained from sequencing populations of the CRISPR libraries cultured in the presence or absence of Golgicide A (GCA), a potent and specific inhibitor of the GEF activity of GBF1 (Sáenz et al., 2009) (Fig. 2A). The concentration of 4 µM GCA used in the screen was determined prior to the screen to maximize both enrichment, i.e., positive selection for rescue of compound toxicity, and depletion, i.e., negative selection for SL interactions (Fig. S1). sgRNA frequencies were determined by sequencing, and relative fold changes in sgRNA abundances between GCA- and mock-treated samples were reported (Table S1 and Fig. 2B).

**Figure 2.**
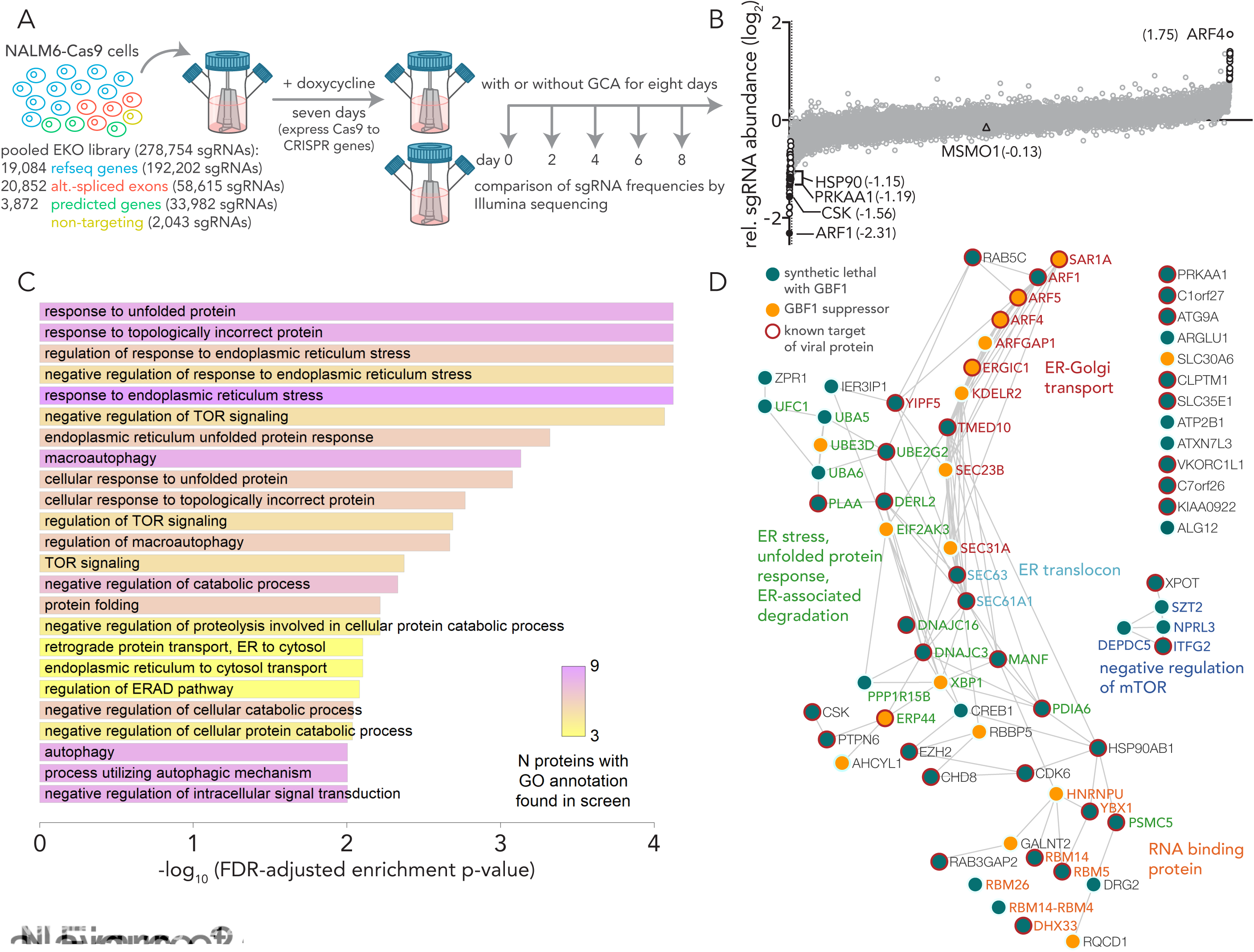
A chemogenomic screen identifies synthetic lethal partners of GBF1. (**A**) A schematic of the experimental design for chemogenomic screening with the GBF1 inhibitor golgicide A (GCA). A CRISPR extended knockout (EKO) library of NALM6-Cas9 cells was treated with 2 µg/ml doxycycline to induce individual gene knockouts via Cas9 expression. The pooled library was split into individual flasks and grown over 8 days period in the presence or absence of 4 µM golgicide A (GCA). Following incubation, guide RNA frequencies were measured using Illumina sequencing, and log_2_ fold changes between GCA and control samples were compared. (**B**) A plot of relative sgRNA frequencies of all genes showing genes passing a 0.05 FDR cutoff in white circles. The 53 genes with negative sgRNA fold change from GCA treatment represent putative synthetic lethal interactors of *GBF1*. The 17 genes with overrepresented sgRNAs and positive sgRNA fold change represent *GBF1* suppressors that may confer protection against GCA. (**C**) Gene ontology functional enrichment analysis of synthetic lethal partners of *GBF1*. The 53 putative synthetic lethal partners of *GBF1* were analyzed in clusterProfiler against the entire KO gene collection from the CRISPR library to functionally classify the SL genes. Significantly enriched gene ontologies are plotted and ranked by their -log_10_, FDR-adjusted enrichment p-value. The number of putative synthetic lethal genes in each gene ontology is coded by the heatmap and ranges from 3 (yellow) to 9 (pink). (**D**) A combined PPI network of the 53 synthetic lethal interactors of *GBF1* (green circles) and the 17 *GBF1* suppressors (orange circles) was obtained from the STRING database and visualized using Cytoscape. Edges between two circles denote evidence-based interaction between the connecting proteins. Circles with red outlines highlight known targets of viral proteins, as per the VirHostNet (v2.0) virus-host PPIs database. Gene names of the proteins and their gene ontology functions are color matched.

Using an FDR cutoff of <0.05, there were 53 underrepresented genes and 17 overrepresented genes in the GCA treated samples relative to the controls (Fig. 2B; white circles). Underrepresented genes represent putative SL partners of GBF1 and the top SL candidate, ADP-ribosylation factor 1 (ARF1), is a small GTPase that regulates the recruitment and assembly of COP I on Golgi and ERGIC membranes (Liang and Kornfeld, 1997). GBF1 facilitates GDP to GTP exchange on ARF1 to regulate recruitment of effectors such as coat protein and lipid-modifying enzymes to ARF1-localized membrane sites and creates a domain competent for secretory cargo transport (Claude et al., 1999; Donaldson and Jackson, 2011; Kawamoto et al., 2002). In yeast, a negative genetic interaction exists between *ARF1* and the yeast GBF1-ortholog, guanine nucleotide exchange on ARF 1 (*GEA1*) (Surma et al., 2013), and *GEA1* overexpression rescues an *arf1*Δ temperature sensitive growth defect (Chantalat et al., 2003). Functional enrichment analysis of the 53 putative SLs of GBF1 showed enrichment for genes in the early secretory pathway, and genes involved in the misfolded protein-triggered ER stress response (Fig. 2C, D). *GBF1* depletion is known to induce an unfolded protein response in the ER (Citterio et al., 2008). Thus, putative SLs of GBF1 are likely functionally redundant with GBF1, an attribute of the genetic interactions between SLs that offers buffering in the event of a loss of function of one of the SL genes (Hartman IV et al., 2001; Mast et al., 2020). As evident by a PPI network, the 53 *GBF1*-SLs and the 17 *GBF1* suppressors are functionally related and can be grouped into a few distinct functional clusters (Fig. 2D). For example, one cluster of *GBF1*-SLs are enriched in ER stress, unfolded protein response, and ER-associated protein degradation pathways, while eight out of the seventeen *GBF1* suppressors contribute to ER-Golgi vesicular transport (Fig. 2D). GBF1-SLs also include a cluster of RNA binding proteins, and members of the KICSTOR (Wolfson et al., 2017) and DEPTOR (Peterson et al., 2009) complexes that negatively regulate mTOR signaling (Fig. 2D). Several genes from both lists possess GTPase activity, e.g., the *GBF1*-SLs: *ARF1, TMED10, DRG2, RAB5C, YIP5, RAB3GAP2*, and the *GBF1* suppressors: *ARF1GAP1, SAR1A1, ARF4*, and *ARF5* (Fig. 2D). ARF4, a class II ARF implicated in endosomal morphology and retrograde transport to the Golgi (Nakai et al., 2013), was the top overrepresented gene (Fig. 2B). A previous large-scale insertion mutagenesis screen identified a role for ARF4 in conferring resistance to Golgi disrupting agents (Reiling et al., 2013), suggesting a protective role of ARF4 against GCA toxicity. Just over half of the GBF1-SLs and suppressors identified in our screen are directly targeted by viral proteins (Fig. 2D, circles with red boundaries).

We searched this list against the Drug Gene Interaction Database (Cotto et al., 2018) for potential ‘druggability’ (http://dgidb.org/search_categories) and selected four druggable, putative synthetic lethal interactors of *GBF1* (Fig. 2A). In addition to *ARF1*, we selected: heat-shock protein 90 (*HSP90*), a protein chaperone with ATPase activity (Rowlands et al., 2010); C-terminal Src kinase (*CSK*), which negatively regulates Src family kinases and has roles in cell growth, differentiation, migration and immune response (Okada, 2012); and protein kinase, AMP-activated, alpha 1 (*PRKAA1*), the catalytic subunit of the 5’-prime-AMP-activated protein kinase (AMPK) with roles in regulating cell stress and metabolism (Sanli et al., 2014). These four putative SL partners of GBF1 were silenced in HeLa cells along with methylsterol monooxygenase 1 (*MSMO1*), included as a control because it did not show depletion or enrichment in sgRNA abundance, and *ARF4*, because it was the topmost significantly overrepresented in our drug CRISPR screen (Table S1 and Fig. 2A). The knockdown (KD) cell lines were incubated with 1.5 µM or 4 µM of GCA/DMSO-alone for 48 h before measuring viabilities. Synthetic lethal effects of combining GCA with shRNA-mediated depletion were observed in *ARF1* KD cells with only 40% or 50% viability as compared to the DMSO-treated cells at 4 µM and 1.5 µM GCA concentration, respectively (Fig. 3A). When the viability of each KD cell line was compared to that of the *MSMO1* KD control, the decrease in the viability was statistically significant for *ARF1* KD at both concentrations, confirming the results of the chemogenomic screen. A GCA dose-response assay monitoring cell growth inhibition as a function of GCA concentration showed a nearly two-fold reduction in IC_50_ value for *ARF1* KD cells as compared to the control (Fig. 3B), further validating the SL interaction between *GBF1* and *ARF1*.

**Figure 3.**
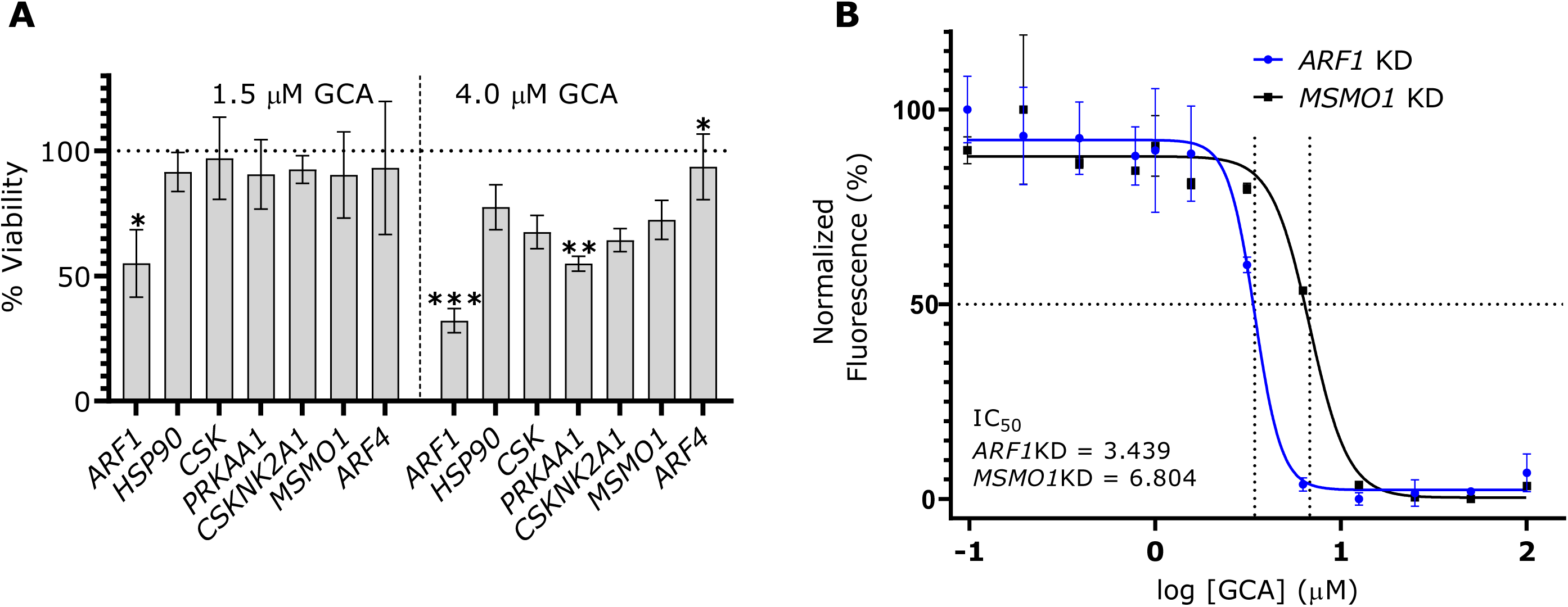
Validation of putative synthetic lethal interactions in HeLa cells. (**A**) *ARF1* displays a robust synthetic lethal interaction with *GBF1. ARF1, HSP90, CSK, PRKAA1*, the control gene *MSMO1*, and the top *GBF1* suppressor gene *ARF4* were silenced in HeLa cells with shRNA-mediated lentivirus transductions and incubated with 1.5 or 4 µM golgicide A (GCA) or DMSO for 48 h. CellTiterBlue reagent was added and fluorescence measurements were collected. Live cells metabolize the reagent into fluorescent products and an increase in the fluorescence signal is directly proportional to the number of living cells. The percent viability at each GCA concentration was calculated by dividing the fluorescence from a GCA-treated sample by its matched DMSO-treated control. Changes in cell viabilities for each knockdown (KD) cell line were determined by comparing the respective percent viabilities to the *MSMO1* KD control using a Brown Forsythe and Welch ANOVA multiple comparison test (Brown and Forsythe, 1974; Welch, 1951), with statistically significant differences are indicated as: * if *p*-value < 0.01; ** if *p*-value < 0.001; *** if *p*-value <0.0001. Error bars represent the SEM from three biological replicates. (**B**) *ARF1* KD cells show enhanced sensitivity in a GCA dose-response curve. A GCA or DMSO working solution (200 µM) was serially diluted and co-plated with 20,000 cells per well of *ARF1* KD and *MSMO1* KD cells in a 96-well plate, with final GCA or DMSO concentrations ranging from 0 – 100 µM. After 48 h, cell viability was measured with CellTiterBlue and the normalized fluorescence, relative to the DMSO-treated samples, was calculated using the smallest and largest mean values to define 0% and 100%, respectively. A dose reponse curve of the normalized fluorescence was plotted against the log_10_ GCA concentration and IC_50_ values were calculated using the a four parameter logistic regression model in Graphpad. Error bars represent the SEM from three biological replicates.

The premise of SL-driven antivirals is that a viral infection disrupts normal protein functions consequently generating vi-hypomorphs in infected cells. As a result, the infected cells may become more vulnerable to drugs that target SL partners of the vi-hypomorph than in uninfected cells lacking the vi-hypomorph. We tested this hypothesis in the context of expressing poliovirus 3A because it recruits GBF1 to sites of poliovirus replication (Belov et al., 2007, 2008a; Richards et al., 2014). The dynamics of GBF1-3A interactions observed during viral infection, including GBF1-mediated ARF1 activation and translocation, are retained in cells ectopically expressing the viral protein alone (Belov et al., 2005, 2007; Richards et al., 2014; Wessels et al., 2006a), which allows for testing the formation of a GBF1 vi-hypomorph in a simpler yet relevant model system without the confounding effects of viral infection. We tagged poliovirus protein 3A with a modified FLAG epitope (FLAG*) (Teterina et al., 2011) and transiently expressed it in HeLa cells (Fig. 4). Poliovirus isolates expressing a modified FLAG tagged 3A are stable and yielded 3A-tagged viruses with similar fitness to the wild-type untagged virus, suggesting that this protein behaves in a similar manner as the wildtype protein (Teterina et al., 2011). The FLAG* tag is similar to the conventional 8-amino acid (DYKDDDDK) FLAG tag, but with the last aspartic acid replaced by tyrosine and is recognized by α-FLAG antibodies (Teterina et al., 2011) (Fig. 4A). N-terminal, FLAG*-tagged 3A, or an empty plasmid control was transiently transfected into HeLa cells for 24 h. 3A-FLAG* and associated proteins were affinity purified on magnetic beads conjugated with α-FLAG antibodies (Fig. 4A). Isolated complexes were washed, and the eluate, along with a fraction of the load and wash were resolved by SDS-PAGE, transferred to nitrocellulose, and probed with α-FLAG and α-GBF1 antibodies (Fig. 4A). A band of ∼10 kDa was detected in the eluate corresponding to the FLAG*-3A protein and a slower migrating, high molecular weight band of ∼200 kDa, corresponding to GBF1, was detected in the eluate of GBF1 immunoprecipitated from cell expressing 3A-FLAG*, but not from cells transfected with the empty plasmid control (Fig. 4A). This observation confirmed that the ectopically expressed 3A-FLAG* protein retained its ability to physically interact with GBF1, as reported previously (Teterina et al., 2011).

**Figure 4.**
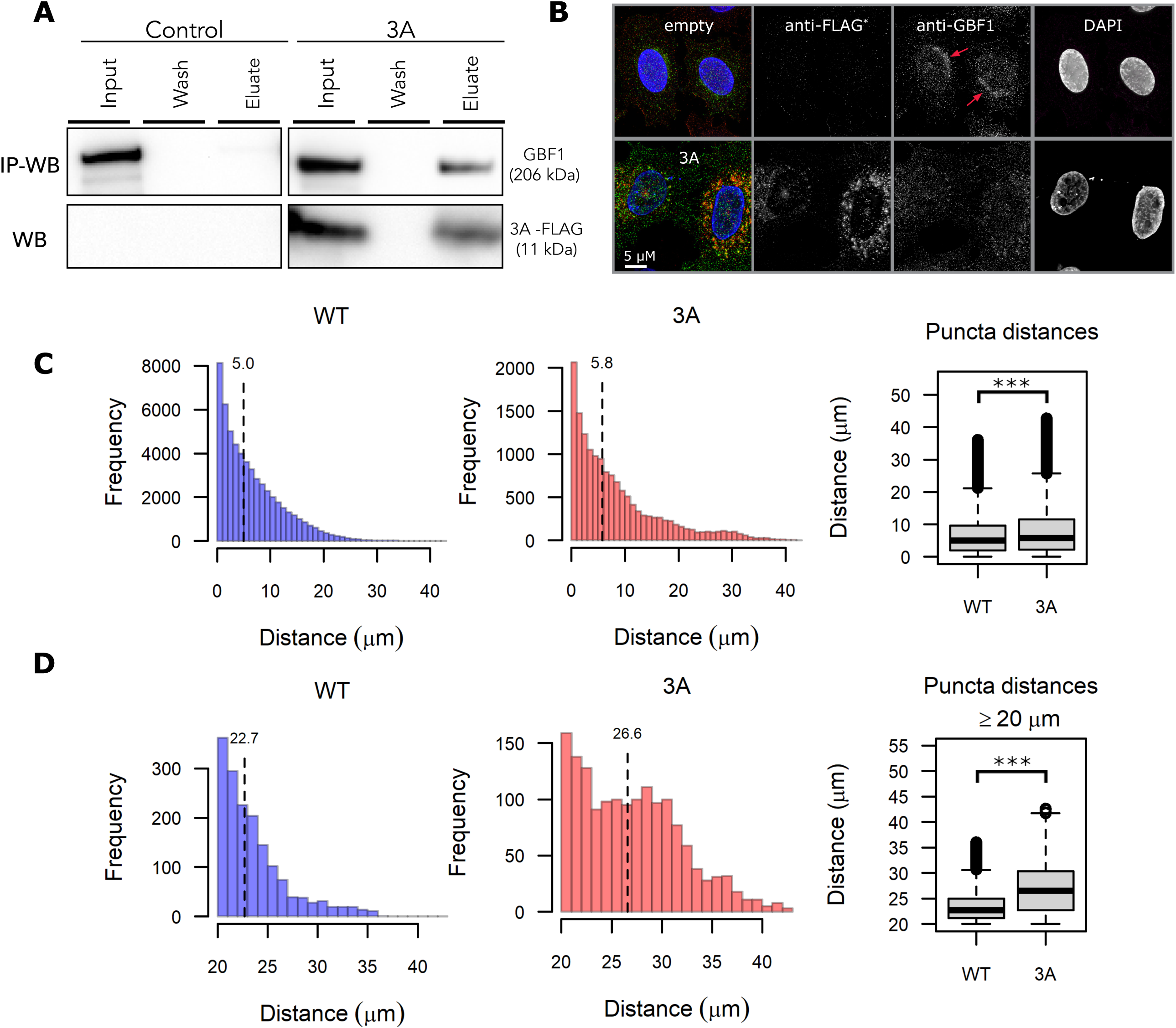
Poliovirus nonstructural protein 3A induces a *vi-hypomorph* of GBF1. (**A**) Poliovirus 3A physically interacts with GBF1. HeLa cells were transfected with FLAG* tagged poliovirus 3A or an empty control plasmid for 24 h. Equal amounts of lysates were prepared, and immunoaffinity enriched for bound protein complexes to 3A-FLAG* protein. Affinity captured proteins were eluted and resolved on SDS-PAGE along with 1% of the total input lysate and the final wash. Resolved proteins were transferred to a nitrocellulose membrane and immunoblotted using anti-GBF1 (top panel) and anti-FLAG (bottom panel) antibodies. (**B**) Poliovirus 3A redistributes GBF1 away from its perinuclear localization. HeLa cells transfected with FLAG* tagged poliovirus 3A or an empty control plasmid were fixed, stained with fluorescently labeled antibodies against FLAG and GBF1, and imaged by wide-field fluorescence microscopy. Bar 5 µM. (**C-D**) Images of GBF1 were analyzed and the distances of each GBF1 puncta to the nearest nucleus was determined and plotted across the entire distance range (C) and between 20 µM to 40 µM (D) for 42 control cells and 13 cells transfected with 3A-FLAG*. The corresponding box plots show statistically significant differences in GBF1 distribution between the two samples with *** representing a *p*-value <0.0001.

We next tested if the physical interaction between 3A and GBF1 had consequences for GBF1 function, suggestive of a GBF1 hypomorph. HeLa cells were transduced with lentivirus delivering an empty control or 3A-FLAG*and cells were fixed and immunostained with α-FLAG-647 (red) and α-GBF1-488 (green), antibodies (Fig. 4B). GBF1 was visualized as puncta enriched in a juxtanuclear position consistent with a Golgi localization in the control cells (Fig. 4B; red arrows). This juxtanuclear enrichment was lost in 3A transduced cells and the GBF1 puncta were instead found redistributed throughout the cytoplasm (Fig. 4B). We quantified this redistribution by measuring the distance of all GBF1 puncta from the nearest nucleus for each cell in the dataset (Fig. 4C, D). This quantification revealed an increase in GBF1 puncta localized away from the nucleus in 3A transduced cells that was statistically significant compared to control. The depletion of Golgi-localized GBF1 upon expression and interaction with poliovirus 3A is consistent with 3A inducing a GBF1-hypomorph.

Having identified *ARF1* as a SL interactor of *GBF1* and established that a GBF1 hypomorph may be induced by expression of poliovirus 3A, we next asked if the 3A-induced hypomorphic state of GBF1 was sufficient to drive a synthetic lethal interaction in cells depleted of ARF1. HeLa cells were treated with shRNA to *ARF1*, or, as a control, shRNA to *MSMO1*, which had no effect in the *GBF1* SL screen (Fig. 2B), and the depletion of ARF1 was evaluated by western blotting (Fig. S2). 3A-FLAG*or an empty plasmid control was transiently transfected into both KD cell lines and expression of 3A-FLAG* was detected by flow cytometry (Fig. 5A). The viability of the *ARF1* KD cells was significantly decreased by the expression of 3A-FLAG* (Fig. 5B). By comparison, the *MSMO1* KD cells were not significantly affected (Fig. 5B). Importantly, this decrease in cell viability, 30% ± 3.5, (Fig. 5B) was comparable to the percent of *ARF1* KD cells expressing 3A-FLAG*, 37% ± 3.0 (Fig. 5A), supporting our hypothesis that the 3A expressing cells in which a hypomorph of GBF1 was generated due to GBF1-3A interaction were selectively killed only in the absence of a functional *ARF1* (Fig. 5).

**Figure 5.**
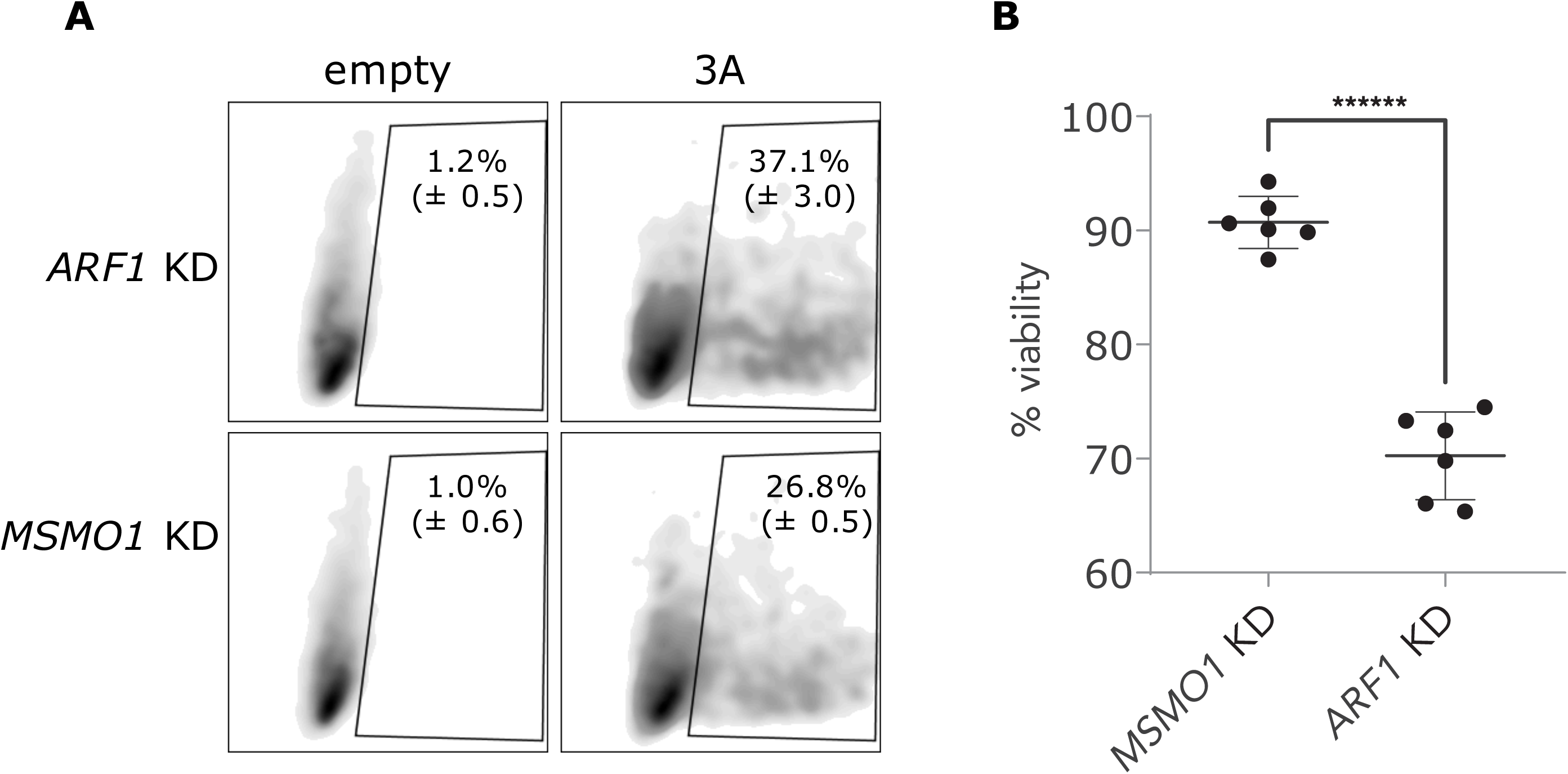
Synthetic lethal killing of a vi-hypomorph of GBF1. (**A**) Expression of poliovirus 3A-FLAG* in *ARF1* and *MSMO1*. Each gene was stably silenced in HeLa cells and transfected with FLAG*-tagged poliovirus 3A. The expression levels of 3A-FLAG 48 h post transfection were measured by flow cytometry. (**B**) Poliovirus 3A induces cell death in *ARF1* KD cells. Cell viabilities of *ARF1* KD and *MSMO1* KD cells transfected with 3A-FLAG* or an empty plasmid control were measured using the fluorescence readout of CellTiterBlue. A change in fluorescence is directly proportional to the number of living cells, and a decrease in the absolute fluorescence indicates reduced cell viability. Percent viabilities of ARF1 KD and MSMO1 KD cells were calculated by dividing the absolute fluorescence values of 3A-transfected samples by the matched empty-transfected samples. Multiple t-test was used to compare percent viability between the 3A-treated *ARF1* KD and *MSMO1* KD cells with ****** representing P-value < 0.000001. Error bars represent the SEM from six biological replicates.

Synthetic lethality is conventionally described as a type of genetic interaction between two nonessential genes that participate in a parallel or redundant process to carry out an essential function, where mutations in either gene alone does not affect cell viability, but mutations in both genes results in cell death (Nijman, 2011b). In this manuscript, we have extended the idea of synthetic lethality to interactions involving a virus-induced hypomorph. In poliovirus, the viral replication complex protein 3A physically interacts with GBF1 (Belov et al., 2008a; Teterina et al., 2011) and re-localizes it to poliovirus replication complexes (Richards et al., 2014). We show here, that this re-localization sensitizes and reduces the viability of cells depleted of ARF1.

In summary, viral-host protein-protein interactions that result in the functional attenuation of the host factor offer a promising avenue for therapeutics based on the principle of synthetic lethality (Mast et al. 2020). In this context, the infected cell becomes selectively dependent upon the otherwise nonessential SL partner of the vi-induced hypomorph. Considering that GBF1 is a common target of many viruses, it is tempting to speculate that SL interactors of *GBF1*, or other common proviral factors, might be candidates for broad-spectrum host-based antivirals. Our proof-of-concept experiments provide a rational approach for identifying these novel antivirals. Our strategy to target SL interactions of the vi-induced hypomorph has a potential to change the current paradigm for host-based therapeutics that can lead to broad spectrum antivirals and can be applied to other intracellular pathogens.

## Materials and Methods

### Cell culture and plasmids

HeLa cells (ATCC CCL-2) and HEK293-FT (ThermoFisher) were cultured at 37 °C in 5% CO_2_ in medium composed of high glucose Dulbecco’s modified eagles medium (DMEM, Gibco) supplemented with 10% (v/v) heat inactivated fetal bovine serum (VWR), 1× penicillin/streptomycin (ThermoFisher Scientific), 20 mM L-glutamine (Gibco), 1× nonessential amino acids (Gibco), 1× sodium pyruvate (Gibco) and 10 mM HEPES buffer (Gibco) (complete media).

Cell line authentication was provided by the American Type Culture Collection and ThermoFisher Scientific. In general, cells were passaged 5-10 times and periodically tested for contamination using MycoAlert™ Mycoplasma Detection (Lonza) kit.

A doxycycline-inducible Cas9 clonal cell line of NALM-6 cells (Wang et al., 2009) was cultured at 37 °C in 5% CO2 in RPMI-1640 medium supplemented with 10 % (v/v) heat inactivated fetal bovine serum, as described (Bertomeu et al., 2017).

Agarose stabs of *E. coli* (DH10B) harboring custom-made mammalian expression vector pD2109-EF1 was purchased from ATUM (Newark, CA USA). pD2109-EF1 encodes poliovirus protein 3A with a modified FLAG tag (DYKDDD<u>Y</u>K) inserted at the N-terminus. The modified FLAG tag (referred here as FLAG*), contains a tyrosine (Y) residue at position 7 instead of aspartic acid residue (D) found in a typical FLAG tag sequence. A complete sequence of the 3A-FLAG*protein with the inserted FLAG* tag (in bold) is as follows:

GPLQYK**DYKDDDYK**DLKIDIKTSPPPECINDLLQAVDSQEVRDYCEKKGWIVNITSQVQTERNIN RAMTILQAVTTFAAVAGVVYVMYKLFAGHQ

Plasmid DNA was purified from the agarose stabs using NucleoBond Xtra Midiprep kit (Macherey Nagel) by following manufacture’s protocol. 3A-FLAG* protein was expressed in HeLa cells by plasmid DNA transfection. We used an empty plasmid pLKO1.puro with comparable size to pD2109-EF1 as a control for transfections.

Bacterial glycerol stocks of MISSION® shRNAs were purchased from Sigma Aldrich for: *ARF1* (clone ID: TRCN0000039874, TRCN0000039875), *ARF4* (TRCN0000298174, TRCN0000047940), *MSMO1* (TRCN0000230198, TRCN0000046245), CSK(TRCN0000199500, TRCN0000000804), *HSP90* (TRCN0000008747,TRCN0000315415), and *PRKAA1* (TRCN0000000861, TRCN0000000859). shRNA plasmid DNA was purified using NucleoBond Xtra Midiprep kit by following manufacture’s protocol.

### Chemogenomic screening and data analysis

Genome-wide custom extended-knockout (EKO) pooled library was created in a B-cell lymphoma line using a published protocol (Bertomeu et al., 2018). Briefly, a clone of the NALM-6 cells expressing Cas9 under a doxycycline-inducible promoter was transduced with the 278K sgRNAs followed by selection over blasticidin, and induction of knockdown of genes with 2 µg/ml doxycycline over a seven-day period. At that time (day 0), the EKO library was split into separate flasks, one containing 4 µM GCA, three containing media alone and two containing 0.1% DMSO and each library flask was grown for eight more days. During this period, cell counts were made every two days and population doublings were monitored. After each cell count, cells were diluted down to 28 million cells per flask and fresh media was added. Whereas all other samples were grown in T-75 flasks from days 0-8 of the screen, one of the untreated control samples was grown in a T-175 flask and was diluted down to 70 million cells every two days instead of 28 million. sgRNA sequences were recovered by PCR of genomic DNA, reamplified with Illumina adapters, and sequenced on an Illumina HiSeq 2000 instrument. The GCA-treated sample DNA was later re-sequenced on an Illumina Next-Seq 500 instrument to increase coverage. Illumina sequencing reads were aligned to the theoretical EKO library using Bowtie 2.2.5, with the -norc (no reverse complement) option and otherwise default parameters. sgRNA read counts were tabulated from all successfully aligned reads. Having found no significant differences between untreated and 0.1% DMSO-treated controls, we opted to add together the sgRNA read counts from all control samples to generate a more robust estimate of the expected sgRNA frequency distribution. We used RANKS (Robust Analytics and Normalization for Knockout Screens) (Bertomeu et al., 2018) with default parameters to generate gene scores p-value and FDR values, comparing the sgRNA read counts of the GCA-treated sample to those of the controls (Table S1). We also calculated gene-level log_2_ fold-changes in sgRNA representation by first summing across each sample the reads of all (usually 10) sgRNAs targeting the gene to calculate a single ratio normalized to the ratio of total aligned read counts per sample (Table S1). This approach effectively downweighs less well represented guides in contrast to the traditional approach of taking the average of the individual sgRNA fold-changes. Reported gene essentiality and essentiality rank in Table S1 are from a previous screen (Bertomeu et al., 2018).

Ontology biological process enrichment analysis was performed on the putative GBF1-SLs (FDR < 0.05) using ClusterProfiler (Yu et al., 2012) where the list was analyzed against the entire KO genes from the CRISPR library to functionally classify the SL genes.

The list of 70 genes (53 GBF1-SLs and 17 GBF1 suppressors) passing the FDR cut off (< 0.05) was submitted to the STRING database (Szklarczyk et al., 2019) to map evidence-based PPIs with the active interaction sources: textmining, experiments, databases, co-expression, neighborhood, gene-fusion, and co-occurrence. The resulting PPI network was visualized using Cytoscape (3.8.2) (Shannon et al., 2003) in a radial layout, with proteins denoted as circles and interactions as edges. The 70 genes were also searched against a virus-host PPI database, VirHostNet (v2.0) (Navratil et al., 2009), to identify known interactors of virus proteins.

### 10× Lentivirus stock preparation

Glycerol stocks of the validated MISSION shRNA vectors for *ARF1, PRKAA1, HSP90, CSK, MSMO1* and *ARF4*, were obtained from Sigma Aldrich (St. Louis, MO). 10× stocks of non-replicating lentiviral stocks were generated by transfection of HEK293-FT cells as follows: 4 × 10^6^ HEK293-FT cells were plated on poly-L-lysine coated 10 cm dishes to achieve 70-80% confluency at time of transfection. The following day, transfection mixtures were prepared by mixing 20 µl Polyethylenimine MAX (Polysciences Inc, Warrington, PA) prepared at 1 mg/ml, together with 4.75 µg of transgene shRNA constructs, 1.5 µg of viral envelope plasmid (pCMV-VSV-G), and 3.75 µg of viral packaging plasmid (psPax2). After incubating for 10 min at room temperature in DMEM, transfection complexes were added dropwise to cells. After overnight incubation, cells were washed to remove the transfection mixture and replaced with 10 ml of pre-warmed media. Lentivirus-containing supernatant was harvested 48 h later, centrifuged for 5 min at 900 *g* to remove cell debris, passed through 0.45 µm syringe filters, and collected by centrifugation for 4 h at 78,900 *g*. Supernatants were decanted and pelleted lentiviruses were re-suspended in 0.1 ml Opti-MEM (Gibco) to obtain 10× lentivirus concentrates and stored at −80 °C until use. A similar protocol was used to prepare 10× lentivirus stocks of 3A-FLAG*.

### shRNA-mediated gene knockdowns

To induce knockdown of the top putative GBF1 SL genes, 300,000 HeLa cells were transduced with lentiviral supernatants in 6-well plates. At time of plating, 10× lentivirus concentrates were diluted in 1 ml of Opti-MEM containing 8 × 10^−3^ µg/ml of polybrene (Sigma Aldrich; St. Louis, MO) and incubated overnight at 37 °C. The following day, the transfection mix was replaced with 2 ml of pre-warmed complete media and incubated for 24 h. In order to select for cells with stable integration of shRNA transgenes, overnight media was replaced with complete media containing 1.5 µg/ml puromycin. Cells were selected for at least 3 days prior to experiments. Stably silenced knockdown cell lines (*ARF1* KD, *HSP90* KD, *CSK* KD, *PRKAA1* KD, *MSMO1* KD, and *ARF4* KD) were harvested by trypsinization in 0.25% trypsin-EDTA, washed in pre-warmed PBS, cells were counted, and a small aliquot was cells were saved for western blot analysis to verify protein level knockdown efficiencies for each gene.

### Western blot analysis

Cell pellets of the KD cell lines were re-suspended in a chilled IP lysis buffer (20 mM HEPES-KOH, pH 7.5, 1% (w/v) Triton X-100, 0.5% (w/v) sodium deoxycholate, 110 mM potassium acetate, 2 mM MgCl_2_, 25 mM NaCl, and 1× cOmplete protease inhibitor cocktail (Roche)) and lysed by sonicating for 1 min using a probe sonicator (QSonica) operated at an amplitude of 10 with 10 s on-off cycles. Lysates were centrifuged at ∼100,000 *g* for 5 min and supernatants were transferred into a fresh tube. Protein concentrations were determined by bicinchoninic acid assay (ThermoFisher Scientific) and working solutions of lysates at concentrations of 15-30 µg total proteins per 30 µl were prepared with 1× lithium dodecyl sulfate (LDS) sample buffer with reducing reagent (NuPAGE, ThermoFisher Scientific) followed by heating at 70 °C for 20 min on a Thermomixer (Eppendorf). 30 µl of the reduced lysate was loaded per well on 4-20% or 12%, for *ARF1* KD and *ARF4* KD, Bis-Tris gels (NuPAGE, ThermoFisher Scientific) and protein bands were resolved at a constant voltage of 170 V for 1 h. Protein bands were transferred on PVDF membranes using Xcell2 blot module (ThermoFisher) for 2 h at a constant voltage of 37 V and membranes were blocked in 5% (w/v) milk in TBST for 1 h at room temperature. After blocking, membranes were incubated with primary antibodies (Abcam and GeneTex) against proteins of interests: ARF1 (ab58578 at 1:1000 dilution), ARF4 (ab190000 at 1:1000 dilution), HSP90 (GTX101448 at 1:1000 dilution), CSK (GTX107916 at 1:500 dilution), CSNK2A (13-453 at 1:500 dilution), PRKAA1 (ab32047 at 1:1000 dilution) and MSMO1 (ab116650 at 1:500 dilution) and washed thrice in TBST buffer before incubating with appropriate HRP-conjugated secondary antibodies (goat α-mouse or α-rabbit; 1:2500 dilution). Following incubation, membranes were washed and developed using chemiluminescent substrates (Advansta WesterBright). Images were acquired using a FluorChem imager (Protein Simple), and membranes were stripped using a stripping buffer (ThremoFisher), blocked, incubated with HRP-α-β-actin (ab49900; 1:25,000) and imaged as before. Images were cropped, adjusted for brightness and contrast, and labelled using Adobe Photoshop and InDesign.

### FLAG*-tagged 3A lentivirus direct plasmid transfection

FLAG*-tagged 3A plasmid DNA were directly transfected in HeLa cells for performing immunofluorescent imaging and co-immunoaffinity purification assays using TransIT transfection reagent (Mirus) by following the manufacturer’s recommended protocol. Briefly, transfection mix was prepared in a serum free Opti-MEM media by adding 3A-FLAG* DNA and TransIT reagent in a ratio of 1:3 (wt/v) and the mixture was incubated for 30 min at room temperature. After incubation, the transfection mix volume equivalent to 1 µg and 15 µg total DNA was added to 60,000 HeLa cells for immunofluorescence imaging and 3 × 10^6^ cells for co-immunoaffinity purification assays, respectively.

### Flow cytometry

Transduced HeLa cells were trypsinized using 0.05% (w/v) Trypsin-EDTA and transferred into a U-bottom 96-well plate. Cells were washed twice in PBS supplemented with 10% (v/v) FBS and incubated with Live/Dead Fixable stain (excitation/emission: 416/451; ThermoFisher Scientific) for 15 min on ice. Excess stain was removed by washing twice and cells were fixed using 4% (w/v) paraformaldehyde (Sigma) for 30 min. Fixative solvent was removed, and cells were washed thrice in PBS by centrifugation at 700 *g* for 5 min. Supernatant wash solution was removed, and cells were incubated in PBS with 0.1% (w/v) Triton X-100 for 15 min to permeabilize the cells. After cell permeabilization, the cells were blocked for 1 h in PBS containing 2% (w/v) BSA (manufacturer) and 1% (w/v) Triton X-100. Cells were then stained with R-phycoerythrin (PE)-conjugated α-FLAG (637310; Biolegends, 1:1000 dilution) for 1 h on ice. After staining, cells were washed thrice in the blocking buffer and analyzed on a LSRII flow cytometer (BD Biosciences). The percentage of cells expressing 3A-FLAG* were determined by analyzing the flow cytometry data using FlowJo software (Tree Star, Inc.). Cell populations were filtered using the forward and side scatter to remove cell debris and cell doublets. The remaining single cell subpopulation was then divided using the intracellular PE straining into a 3A-FLAG* positive and negative populations and percentages of positive and negative cells of the total single cells were reported. Experiments were performed in duplicate.

### Immunofluorescence microscopy

HeLa cells were plated in 12 well plates containing 12 mm no.1.5 circular glass coverslips (Fisherbrand) at a cell density of 60,000 per well and transfected with FLAG*-3A or an empty plasmid control. Cells were fixed 24 h post-transfection with 2% (w/v) paraformaldehyde (Sigma) for 30 min, permeabilized with 0.1% (w/v) Triton X-100, and blocked with 2% (w/v) BSA, 0.1% (w/v) Triton X-100 in PBS (blocking buffer). After blocking, cells were incubated with rabbit α-GBF1 (abcam; ab86071 1:1000 dilution) and mouse α-FLAG (Sigma F1804; 1:200 dilution) for 1 h followed by 1 h staining with secondary antibodies goat α-rabbit AlexaFluor-488, and goat α-mouse AlexaFluor-594 (Invitrogen), used at a 1/1000 dilution. The coverslips were mounted on Superfrost™ microscope slides (Fisherbrand), nuclei were stained with DAPI, and the cells were cleared with Prolong Glass (Invitrogen). Images were acquired with a 100× NA 1.4 objective (Olympus) on a DeltaVision Elite High-Resolution Microscope (GE Healthcare Life Sciences). Fluorescence excitation was driven by an Insight SSI solid state light engine (Cytiva) and fluorescence emission was collected by a CoolSnap HQ2 CCD camera (Photometrics). The sides of each CCD pixel are 6.45 µm. Image z stacks were acquired with 0.2 µm steps and 25 - 27 images per stack. Images were deconvolved with a classic maximum likelihood estimation algorithm using Huygens software (Scientific Volume Imaging; The Netherlands) and experimentally determined point spread functions captured by imaging PS-Speck™ beads (Invitrogen) under experimental conditions, as done previously (Mast et al., 2018; Vijayan et al., 2019).

### Image quantification

Images were processed using Imaris software (Bitplane) to quantify the number of GBF1 puncta per cell. Initial cell segmentation was performed by summing the fluorescent intensities from all channels and using the ‘Surface’ command to threshold the images. This segmentation was refined using the ‘Cell’ command. Cell nuclei were defined using the DAPI channel and cell boundaries defined using a watershed algorithm seed by the ‘one nucleus per cell’ function in order to split touching cells. Next, the GBF1-488 channel was selected for detecting GBF1 puncta using the ‘detect vesicle’ function. Statistical values for ‘Vesicle intensity sum’ and ‘vesicle distance to closest nucleus’ for each GBF1 puncta per cell were exported for 42 cells from control samples, and from 13 cells of 3A-FLAG* transfected cells. Distances of GBF1 puncta from the nearest nucleus in 3A-FLAG* cells were compared to distances from control cells using Wilcoxon rank-sum tests. Comparisons were made using all observed puncta and, separately, using only puncta more than 20 mm from the closest nucleus.

### Coimmunoprecipitation

HeLa cells were transfected with 1 µg DNA of each of the FLAG*-3A and an empty control plasmid. At 24 h post-transfection, cells were lysed using mild sonication in IP lysis buffer (20 mM HEPES, 1% (w/v) Triton X-100, 2 mM Magnesium chloride, 25 mM Sodium chloride, 110 mM Potassium acetate, and 0.2% (v/v) antifoam B), and clarified by centrifugation at 100,000 *g* for 3 min. Total protein concentrations were measured using BCA assay and 100 µg of lysate from each sample was used in the immunoprecipitation. 8 µg of α-FLAG (F3165, Sigma Aldrich) was conjugated to 10 mg epoxy-coated M-270 magnetic beads (ThermoFisher Scientific) (Cristea and Chait, 2011). The α-FLAG conjugated beads were washed and re-suspended in IP lysis buffer, and 3 mg bead aliquots were added to the clarified lysates. Lysates were incubated with magnetic beads overnight at 4 °C. After three washes with IP lysis buffer, bound proteins were eluted with 50 μl 1× LDS sample buffer and resolved on 4-20% Bis-Tris NuPAGE gel. Additionally, ∼10% of the eluate volume was resolved on a 3-8% Tris-Tricine gel to confirm expression and enrichment of the immunoprecipitated 3A-FLAG* bait. Proteins were transferred to PVDF membranes and immunoblotted with α-GBF1 (ab86071; abcam, at 1:1000 dilution) and HRP conjugated α-FLAG (A8592; Millipore Sigma 1:2000 dilution). The experiment was performed in triplicate.

### Cell viability assay

Cell viability assays were performed in 96-well plates using the CellTiterBlue reagent (Promega). Working concentrations of GCA were prepared by diluting a 10 mM DMSO-solubilized stock in complete media. Equimolar solutions lacking the drug were prepared by diluting neat DMSO. Cells were incubated in 100 µl of GCA or DMSO working solutions for 48 h before 20 µl of the CellTiterBlue reagent was added, and fluorescence was measured 4 h-post addition using a Synergy HTX Multi-mode plate reader (BioTek). The metabolically active cells convert the blue redox reagent into its fluorescent product with the number of live cells directly proportional to the intensity of the fluorescent product. Fluorescence measurements from the drug-treated samples are normalized using the signal from matched DMSO-treated samples.

For the viability assay validating *GBF1*-SL candidates, 20,000 cells of each gene KD cell line (*ARF1* KD, *HSP90AB* KD, *CSK* KD, *PRKAA1* KD, *ARF4* KD, *MSMO1* KD) were seeded per well in 200 µl media. The next day, the media was replaced with 100 µl of GCA or DMSO at 1.5 µg/ml and 4 µg/ml, incubated for 48 h, and cell viability was measured as described above. Normalized cell viabilities of the KD cells were compared to that of the *MSMO1* KD controls.

For generating GCA dose response curves we used an automated high throughput liquid handling system (PipetteMax, Gilson) for co-plating cells and the drug. Stock solutions of GCA or DMSO were prepared in complete media at a concentration of 200 µM, and serially diluted in cell-containing media of *ARF1* KD and *MSMO1* KD (400,000 cells/ml) to obtain 0–100 µM of GCA/DMSO with 20,000 cells/ well, plated in triplicate. Cell viabilities were measured 48 h post-incubation as described above. GCA dose response curves were plotted with GraphPad Prism 8.

For the proof-of-concept SL viability experiments, 60,000 cells of *ARF1* KD and *MSMO1* KD were seeded in a 6-well plate and transfected the following day with 3A-FLAG* or an empty plasmid control. After 24 h post transfection, cells were harvested, counted, and plated in 96-well plate at a density of 5000 cells per well. The remaining cells were processed for flow cytometry analysis to measure transfection efficiencies. Cell viabilities were measured 48 h post-transfection using CellTiterBlue reagent, as described above. Absolute cell viabilities of *ARF1* KD and *MSMO1* KD cells were compared using a multiple T-test, i.e., the two-stage step-up method of Benjamini, Krieser and Yekutieli (Benjamini et al., 2006), in GraphPad Prism 8. The experiment was performed in triplicate.

### Online supplemental material

Table S1 contains supporting data reporting the GCA CRISPR screen results for the 19,029 Refseq genes. For each gene, we report the RANKS (Robust Analytics and Normalization for Knockout Screens) score, associated p-values, the FDR, the number sgRNA considered for the analysis, and the gene-level log_2_ fold changes. Fig. S1 contains the results from a GCA dose-response assay used to determine the concentration of GCA in the CRISPR screen reported in Fig 2. Fig. S2 contains supporting immunoblots that show the efficiency of the shRNA-mediated protein depletion for the *ARF1, MSMO1, ARF4, PRKAA1, CSK*, and *HSP90* KD cell lines.

## Acknowledgments

This work was supported by National Institutes of Health grants R01 GM112108 and P41 GM109824 to J.D. Aitchison, and R21 AI151344 to A. Kaushansky and foundation grant FDN-167277 from the Canadian Institutes of Health Research to M. Tyers/

J.D. Aitchison, A.T. Navare, F.D. Mast, M. Tyers, T. Bertomeu, P. Olivier, and M. Neal are inventors on a provisional patent disclosing synthetic lethal approaches described in this manuscript.

## Author Contributions

J.D. Aitchison, F.D. Mast, and A.T. Navare wrote the original draft of the manuscript. F.D Mast, A.T. Navare, J.P. Olivier, L.N. Carpp, T. Bertomeu, and J. Coulombe-Huntington designed and performed the experiments and interpreted results. M. Neal analyzed data and interpreted results. J.D. Aitchison, A. Kaushansky, and M. Tyers designed and oversaw experiments and revised manuscript drafts. All authors read and commented on the manuscript.

