## Supplemental Materials for "Viral protein engagement of GBF1 induces host cell vulnerability through synthetic lethality"

**Online supplemental material**

Table S1 contains supporting data reporting the GCA CRISPR screen results for the 19,029 Refseq genes. For each gene, we report the RANKS (Robust Analytics and Normalization for Knockout Screens) score, associated p-values, the FDR, the number sgRNA considered for the analysis, and the gene-level $\log_2$ fold changes.

Figure S1

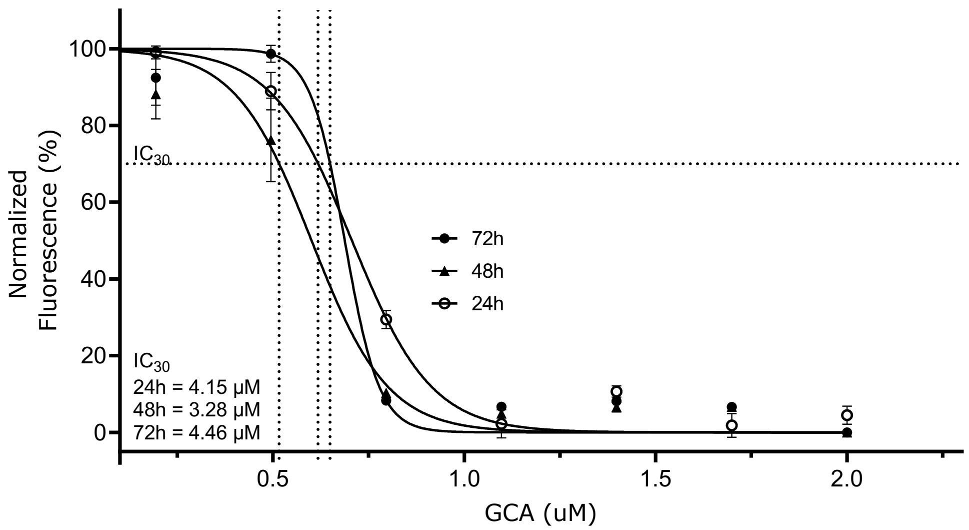


Figure S1. **GCA dose-response assay.** Nalm6 cells were incubated with serially diluted GCA or DMSO, in triplicate. After 24, 48, or 72 h, cellTiterBlue reagent was added to each well and the cells were incubated for 4 h. Metabolically active cells convert the reagent into a fluorescent product, and the fluorescence intensity recorded by a plate reader is directly proportional to the number of live cells. The fluorescence of the GCA-treated samples was normalized to the equivalent DMSO-treated controls and $IC_{30}$ values were determined using the default nonlinear regression model in GraphPad Prism 8. $IC_{30}$ values over the three-day period were averaged ($IC_{30}$ average = ~4.0 µM), to determine the concentration of GCA to be used in the chemogenomic drug screening assay.

Figure S2

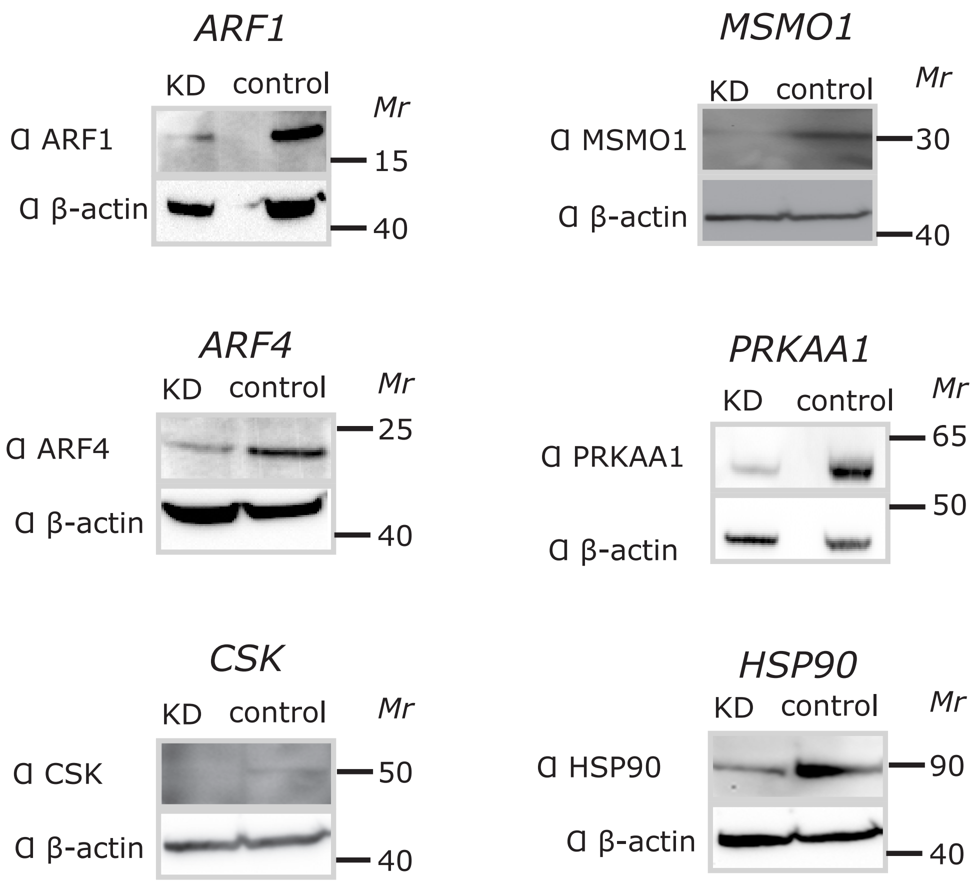


Figure S2. **Validation of shRNA-mediated knockdown.** Four druggable putative synthetic lethal partners of GBF1: *ARF1, PRKAA1*, *CSK*, and *HSP90*; a putative GBF1 suppressor, *ARF4*; and a gene showing no GCA-induced chemogenomic interactions with *GBF1*, *MSMO1*, were silenced in HeLa cells via shRNA-mediated lentivirus transductions and selected on puromycin for 72 h. Small aliquots of a cell pellet from each knockdown cell line were collected, lysed, and the total protein concentrations were measured using a bicinchoninic acid assay. An equal amount of total proteins from the control and knockdown cells were resolved by SDS-PAGE, transferred to nitrocellulose and the presence of each protein of interest was then probed using appropriate antibodies. For the loading control, membranes were stripped, re-blocked and probed using antibodies to β-actin.
